# Temporal Effects on Radiation Responses in Nonhuman Primates: Identification of Biofluid Small Molecule Signatures by Gas Chromatography – Mass Spectrometry Metabolomics

**DOI:** 10.1101/620526

**Authors:** Evan L. Pannkuk, Evagelia C. Laiakis, Michael Girgis, Sarah E. Dowd, Suraj Dhungana, Denise Nishita, Kim Bujold, James Bakke, Janet Gahagen, Simon Authier, Polly Chang, Albert J. Fornace

## Abstract

Whole body exposure to ionizing radiation (IR) (> 0.7 Gy) damages tissues leading to a range of physical symptoms contributing to acute radiation syndrome (ARS). Radiation biodosimetry aims to determine characteristic early biomarkers indicative of radiation exposure (generally at doses > 2 Gy) and is a necessity for effective triage in the event of an unanticipated radiological incident and emergency preparedness. Radiation metabolomics can address this aim by assessing metabolic perturbations following various emergency scenarios (e.g., elapsed time to medical care, absorbed dose, combined injury). Gas chromatography – mass spectrometry (GC-MS) is a standardized platform ideal for chromatographic separation, identification, and quantification of metabolites to discriminate molecular signatures that can be utilized in assessing radiation injury. We performed GC time-of-flight (TOF) MS for global profiling of nonhuman primate (NHP) urine and serum samples up to 60 d after a single 4 Gy γ-ray total body exposure. Multivariate statistical analysis showed a higher separation of groups from urine signatures vs. serum signatures. We identified biofluid markers involved in amino acid, lipid, purine, and serotonin metabolism, some of which may indicate host microbiome dysbiosis. Sex differences were observed amino acid fold changes in serum samples. Additionally, we explored mitochondrial dysfunction by analysis of tricarboxylic acid (TCA) intermediates with a GC tandem quadrupole (TQ) MS platform in samples collected in a time course during the first week (1, 3, 5, and 7 d) after exposure. By adding this temporal component to our previous work exploring dose effects at a single time point of 7 d, we observed the highest fold changes occurring at 3 d, returning closer to basal levels by 7 d. These results emphasize the utility of both MS-based metabolomics for biodosimetry and complementary analytical platforms for increased metabolome coverage.

## 1. Introduction

Exposures to external ionizing radiation (IR) can include occupational hazards [1] or accidents from nuclear energy (e.g., deposition of ^137^Cs after the Chernobyl accident) [2] to normal background radiation [3]. The lingering health risks associated with IR exposures among large populations can be observed in atomic bomb survivors many decades after the initial exposure, both in Hiroshima and Nagasaki [4,5]. More recently, the potential of malicious use of radioactive materials, such as a radiological dispersal device (RDD) or improvised nuclear device (IND), for terrorist actions has been of concern. As such, there is a need for predictive biomarkers indicative of radiation exposure to aid in the triage of large populations during a potential radiological terrorism act. The National Institute of Allergy and Infectious Diseases (NIAID) is leading this effort through the Centers for Medical Countermeasures Against Radiation Consortium (CMCRC) program [6]. Of these needs, our group has particular interest in the use of mass spectrometry (MS) platforms for radiation biomarker development, aiding in rapid high-throughput biodosimetry and assessment of acute radiation syndrome (ARS) and associated tissue damage and recovery [7].

Metabolomics is the collective analysis of small molecules (< 1 kDa) in biofluids, cells, or tissues that provides phenotypic signatures downstream of other –omic technologies, such as transcriptomics or proteomics [8]. By far, the two dominating platforms for metabolomics have been MS and nuclear magnetic resonance (NMR), and as each possesses unique analytical advantages and disadvantages, these platforms are viewed as complementary for increased metabolite coverage [9]. The advantages of NMR platforms include non-destructive sample analysis, high reproducibility, and ability to perform accurate quantification, however, low sensitivity leads to identification of similar profiles of high abundance metabolites and NMR analysis requires large volumes of material [10]. Furthermore, peak deconvolution can be challenging especially in highly complex spectra. MS platforms are capable of producing highly quantitative metabolite concentrations (e.g., tandem quadrupole [TQ] MS), high mass accuracy and resolution (e.g., Fourier-transform ion cyclotron resonance [FTICR] MS), and can provide spatially resolved metabolite profiles when coupled with ambient ionization (e.g., desorption electrospray ionization [DESI]). Increased metabolite coverage and background reduction is achieved through up front separations, such as ultra-performance liquid chromatography (UPLC), gas chromatography (GC), and capillary electrophoresis (CE) [11]. Of the above mentioned, GC-MS platforms have served as the earlier tools in metabolite profiling [12,13], and have become one of the most standardized and reproducible tools for metabolomic profiling [14]. In addition to its superior separation of isomers and small volatile compounds, the breadth of compounds available in reference databases far surpasses other metabolomic platforms [15]. As previous studies on radiation biofluid metabolomics using GC-MS platforms have been limited, including mice (plasma after low dose radiation exposure) [16], rats (urine time effects [17] and time/dose effects [18]; serum dose effects [19]), nonhuman primates (NHPs) (dose effects in urine and serum [20]), and patients (serum after cumulative doses during radiotherapy [21,22]), we have expanded on the temporal effects of radiation exposures on NHP urine and serum for biomarker discovery.

This study examines perturbations to urinary and serum metabolic signatures in NHPs after a 4 Gy γ-ray total body irradiation (TBI) exposure (LD_50/60_ ∼6.6 Gy). This dose was chosen to allow assessment of biofluid signatures over a longer time span without high mortality while representing a potentially lethal level in humans. NHP models are the most relevant animal model for these studies considering they are well characterized in terms of ARS effects, such as hematopoietic and gastrointestinal syndromes and delayed effects, and are more genetically similar to humans than other models [23]. We utilized a GC time-of-flight (TOF) MS platform to globally profile biofluids spanning a time course from pre-exposure to 60 d in an initial discovery phase approach. Identified metabolites implicate perturbation to amino acid, purine, and lipid metabolism. As mitochondrial dysfunction is a known consequence of IR exposure, we refined further our analysis by utilizing a GC-TQ-MS platform to determine fold changes in tricarboxylic acid (TCA) cycle intermediates from pre-exposure to 7 d, showing highest fold changes occurring at 3 d. These results highlight the use of metabolomics to define cellular responses to radiation injury as well as the importance of using multiple analytical platforms to obtain a more complete view of the metabolome.

## 2. Materials and Methods

### 2.1. NHP system

Rhesus monkeys (*Macaca mulatta*) were irradiated at an AAALAC accredited facility as previously described [24]. Briefly, 8 NHPs (4 male, 4 female, average age 6 yrs. and average weight 8.3 kg) were fed commercial chow (twice daily) and fresh fruits, vegetables, or juice (intermittently) and housed in an environment-controlled facility (temperature, 21°C ± 3°C; relative humidity, 50% ± 20%; 12 hr light/dark cycles). NHPs were housed individually ≥ 7 d post-exposure then paired or group housed if deemed healthy. Before irradiation, NHPs were fasted, administered ondansetron (2 mg/ml, 1.5 mg/kg, intramuscular injection) 45 – 90 min before irradiation, and acclimated to procedures related to radiation before exposure to a total dose of 4 Gy at 0.6 Gy/min with a ^60^Co γ source (∼7 min duration, half antero-posterior and half postero-anterior position, doses confirmed with a Farmer^®^ ionization chamber and secondarily with two nanoDot(tm) dosimeters placed on the animals’ body). Animals were again administered ondansetron 30 – 45 min after irradiation for potential emesis. Clinical signs (daily after irradiation) and detailed examinations (1, 3, 5, 7 d and then weekly) were recorded. Pre-exposure (−8 and -3 d) and post-exposure samples (1, 3, 5, 7, 15, 21, 28, and 60 d) were collected, frozen (−80°C), and shipped to Georgetown University Medical Center (GUMC). A previous cohort used for 7 d TCA cycle intermediate and amino acid comparisons has been described in detail [25,26].

### 2.2. Sample preparation and GC-MS instrumentation

#### 2.2.1. Chemicals

Fisher Optima™ grade (Fisher Scientific, Hanover Park, IL, USA) solvents were used for sample preparation. Internal standards were obtained from Cambridge Isotope Laboratories, Inc. (Andover, MA, USA) (citric acid-d_4_, _DL_-glutamic acid-d_5_, _D_-sorbitol-^13^C_1_, and _L_-leucine-d_3_) or Sigma-Aldrich (St. Louis, MO, USA) (4-nitrobenzoic acid). Standards for obtaining retention indices (C_4_-C_24_ FAMEs, methyl nonanoate, and alkane standard mix [C_10_-C_40_]) and for urine pretreatment (urease from *Canavalia ensiformis* [Jack bean] type III) were obtained from Sigma-Aldrich. Derivatization chemicals were obtained from Thermo Scientific (Waltham, MA, USA) (Methoxamine [MOX] reagent and *N*-methyl-*N*-[trimethylsilyl]trifluoroacetamide [MSTFA] with 1% [vol/vol] trimethylchlorosilane [TMCS]).

#### 2.2.3. Sample Preparation

For global metabolomics, urine (60 µl) was aliquoted and treated with 5 µl (160 mg/ml) urease for 1 hr at 37°C with gentle agitation and serum (30 µl) was aliquoted without pretreatment. Quality control (QC) samples were prepared by aliquoting an equal volume from each sample. Samples were deproteinated with 1 ml 100% cold methanol with internal standards (final concentration 20 µg/ml), incubated on ice for 10 min, and centrifuged at 13,000 rpm for 10 min at 4°C. The supernatant was evaporated under N_2_ to ∼100 µl, transferred to a GC vial with a 250 µl glass insert, and evaporated to dryness in a speedvac without heat. Samples were derivatized inline with a Gerstel (Linthicum, MD, USA) multipurpose sampler for 1 hr at 60°C with rigorous shaking with MOX (10 µl) and an additional 1 hr at 60°C with MSTFA / 1% TMCS (90 µl). For TCA intermediate analysis, urine (40 µl) was prepared as above, however, samples were deproteinated with cold methanol (internal standard citric acid-d_4_ at a final concentration of 2 µg/ml) without urease pretreatment and batch derivatized per day.

#### 2.2.4. GC-MS Instrumentation

For global profiling, samples (1.5 µl) were injected into an Agilent (Santa Clara, CA) 7890B GC system (equipped with a Rtx^®^-5 [G27] column [5% diphenyl / 95% dimethyl polysiloxane, 30 m x 0.25 mm x 0.25 µm, 5 m Integra-Guard^®^ column]) mounted to a Leco (St. Joseph, MI) Pegasus HT TOF-MS. The GC settings were as previously described [20,21]: inlet (220°C), transfer line (270°C), oven temperature program (70°C [0.2 min], 70°C – 270°C [10°C/min, held 4 min], 270°C – 320°C [20°C/min, held 2 min]), 1:10 split, carrier gas helium (1.2 ml/min constant gas flow rate), and 220 sec solvent delay. The MS settings and injection program (MAESTRO software) were also as previously described [20,21]: scan range (*m/z* 40 – 600), ion source (200°C), 30 spectra/sec acquisition rate, and liner exchange every 10 samples.

For TCA intermediate analysis, samples were injected into a Waters (Waters, Milford, MA, USA) Xevo™ TQ-GC MS with identical settings as above and data was acquired in multiple reaction monitoring (MRM) mode for pyruvic, citric, isocitric, *cis*-aconitic, α-ketoglutaric (oxoglutaric), malic, succinic, and fumaric acid (Table 1).

**Table 1.** Multiple reaction monitoring transitions and parameters for pyruvic acid, TCA cycle intermediates, and deuterated citric acid (internal standard).

| **Metabolite** | **Parent ion**<br>**(*m/z*)** | **Daughter ion**<br>**(*m/z*)** | **Collision**<br>**(eV)** | **Retention**<br>**time (min)** |
| --- | --- | --- | --- | --- |
| Pyruvic acid | 174 | 74.1 | 20 | 4.69 |
| Citric acid | 273 | 73.1 | 15 | 14.27 |
| Citric acid-d<sub>4</sub> | 276 | 185 | 15 | 14.25 |
| Isocitric acid | 245 | 73.1 | 20 | 14.28 |
| *cis*-Aconitic acid | 229 | 147.1 | 15 | 13.41 |
| α-Ketoglutaric acid | 198 | 73.1 | 20 | 11.48 |
| Malic acid | 233 | 73.1 | 15 | 10.46 |
| Succinic acid | 247 | 147 | 15 | 8.15 |
| Fumaric acid | 245 | 73.1 | 20 | 8.56 |

#### 2.2.5. Data Processing and Statistical Analysis

For global analysis, data preprocessing, deconvolution, and alignment were performed with the software ChromaTOF^®^ v. 4.72.0.0 (Leco, St. Joseph, MI, USA) and the statistical compare function as previously described [20]. Compounds were identified by matching their unique electron ionization (EI) impact spectra to the National Institute of Standards and Technology (NIST) spectral library v. 14 (Gaithersburg, MA, USA) and retention indices to FiehnLib [14]. Pre-exposure (−8 and −3 d) samples were averaged for the control reference. Urine and serum were normalized to an internal standard and urine was further normalized to creatinine. For multivariate analysis, data matrices were imported into MetaboAnalyst 4.0 with K-nearest neighbor missing value imputation, log transformation, and Pareto scaled for visualization by partial least squares-discriminant analysis (PLS-DA) [27]. The performance of the PLS-DA model was assessed using 10-fold cross-validation (CV) and B/W ratio after 1000 permutations [28]. For univariate analyses, data matrices were initially screened for statistically significant compounds with an ANOVA using the proc glm function and a post-hoc Duncan test in SAS 9.4 (Cary, NC, USA). Significant compounds were graphed with GraphPad Prism 6.0 (GraphPad Software, Inc., La Jolla, CA, USA) with outliers removed using robust regression and outlier removal and analyzed with a Kruskal-Wallis test and pot-hoc Dunn’s test across days and a t-test for sex differences. For analysis of TCA cycle intermediates, raw data files were processed and analyzed using TargetLynx v4.1 (Waters, Milford, MA, USA) and log fold changes were graphed with GraphPad Prism 6.0.

## 3. Results and Discussion

Data matrices consisted of 408 compounds for urine and 112 compounds for serum. Creatinine was not significantly different among groups and was used for normalization of urine data. Multivariate PLS-DA analysis was performed to assess separation among pre-IR vs. 1 – 7 d groups and pre-IR vs. 15 – 60 d groups for urine and serum. After potential radiologic emergencies, a 1 – 7 d time course represents a relevant period for initial triage, while at 15 – 60 d more advanced symptoms is expected to have developed but observation and medical treatment will still be required. From 1 – 7 d after irradiation, the 3 d group was the most unique in urine being separated from the pre-IR and 1 d group and 5 – 7 d along component 1 (10-fold cross-validation [CV]: accuracy = 0.67, *R*^2^ = 0.99, *Q*^2^ = 0.80; performance measure: *Q*^2^ P = 0.002; Figure 1A). From 15 – 60 d, the 15 d group separated well along component 1 but higher overlap is seen from 21 – 60 d (10-fold CV: accuracy = 0.44, *R*^2^ = 0.98, *Q*^2^ = 0.65; performance measure: *Q*^2^ P < 0.001; Figure 1B). Separation in serum biosignatures was less pronounced as reflected by their validation and performance measures. Separation along component 1 from 1 – 7 d but higher overlap among groups was evident (10-fold CV: accuracy = 0.40, *R*^2^ = 0.95, *Q*^2^ = 0.57; performance measure: *Q*^2^ P = 0.007; Figure 1C), while little separation was observed from 15 – 60 d (10-fold CV: accuracy = 0.42, *R*^2^ = 0.92, *Q*^2^ = 0.08; performance measure: *Q*^2^ P = 0.009; Figure 1D). While urine metabolite biosignatures are typically more “striking” than serum, as observed in this study, serum is a valuable biofluid when monitoring changes in lipid content after IR exposures and can discriminate exposed individuals from the unexposed at the metabolite level.

**Figure 1.**
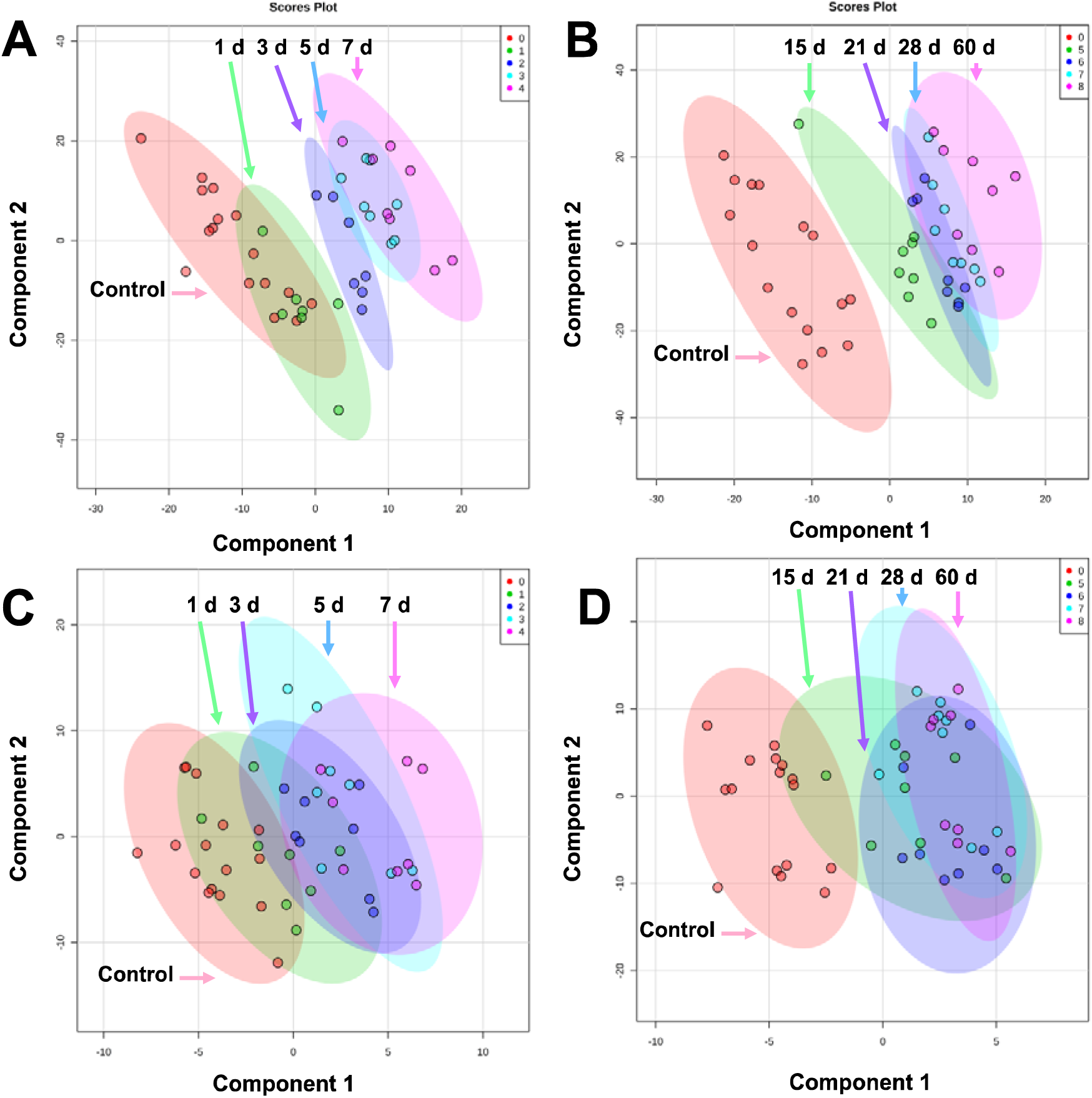
PLS-DA plots comparing pre-exposure to A) 1 – 7 d in urine, B) 15 – 60 d in urine, C) 1 – 7 d in serum, and D) 15 – 60 d in serum after 4 Gy γ-ray TBI in NHPs. Overall, urine showed better separation among groups than serum. The highest separation for both biofluids occur within 15 d and higher overlap is observed from 21 – 60 d (graphs generated in MetaboAnalyst 4.0, pre-exposure samples [-8 and -3 d] were averaged for the control group)

Univariate analyses showed significantly decreased levels of allantoic acid (P < 0.001) from 5 – 60 d and higher levels of 5-hydroxyindoleacetic acid (P = 0.003) at 60 d in urine (Table 2, Figure 2A, Table S1). Allantoic acid is a metabolite produced through purine metabolism downstream of uric acid. Uric acid is enzymatically degraded or spontaneously converted to allantoin, which may be further metabolized to allantoic acid through allantoinase [29]. As Old World primates lack allantoinase (e.g., Rhesus macaques), it is possible that the presence of allantoic acid is due to microbial activity [30,31]. The compound 5-hydroxyindoleacetic acid is a metabolite of serotonin (5-hydroxytryptamine) formed by aldehyde dehydrogenase via 5-hydroxyindoleacetaldehyde and as a possible downstream product of tryptophan metabolism may also be influenced by host microbiota [32,33]. Serotonin has previously received interest as a radioprotector [34,35] and elevated levels of 5-hydroxyindoleacetic acid has been shown in urine after radiation exposure [36–38]. Interestingly, kynurenic acid and xanthurenic acid are also metabolic products derived from tryptophan, possible indicators of renal damage, and are commonly detected in urine, as identified in previous radiation metabolomics studies [7]. Other metabolites in the tryptophan pathway due to host microbiota dysbiosis have been identified in plasma and future studies may continue to expand on these [39].

**Table 2.** Compounds detected by GC-TOF-MS global profiling of NHP biofluids after 4 Gy γ radiation exposure.

| **Biofluid** | **Metabolite** | **Retention index** | **Unique mass** | **P value** | **No. TMS** | **HMDB ID** |
| --- | --- | --- | --- | --- | --- | --- |
| Urine | Allantoic acid | 662102 | 331 | 0.020 | 1 | HMDB 01209 |
|  | 5-Hydroxyindoleacetic acid | 786849 | 290 | 0.003 | 3 | HMDB 00763 |
| Serum | Oleic acid | 787976 | 117 | <0.001 | 1 | HMDB 00207 |
|  | Inosine | 906197 | 73 | 0.002 | 4 | HMDB 00195 |
|  | Leucine | 353596 | 158 | 0.001 | 2 | HMDB 00687 |
|  | Isoleucine | 366901 | 158 | 0.005 | 2 | HMDB 00172 |
|  | Valine | 320280 | 144 | 0.002 | 2 | HMDB 00883 |
|  | Serine | 405133 | 73 | 0.012 | 3 | HMDB 00187 |
|  | Threonine | 420194 | 73 | 0.003 | 3 | HMDB 00167 |
|  | Phenylalanine | 542089 | 73 | 0.004 | 2 | HMDB 00159 |

**Figure 2.**
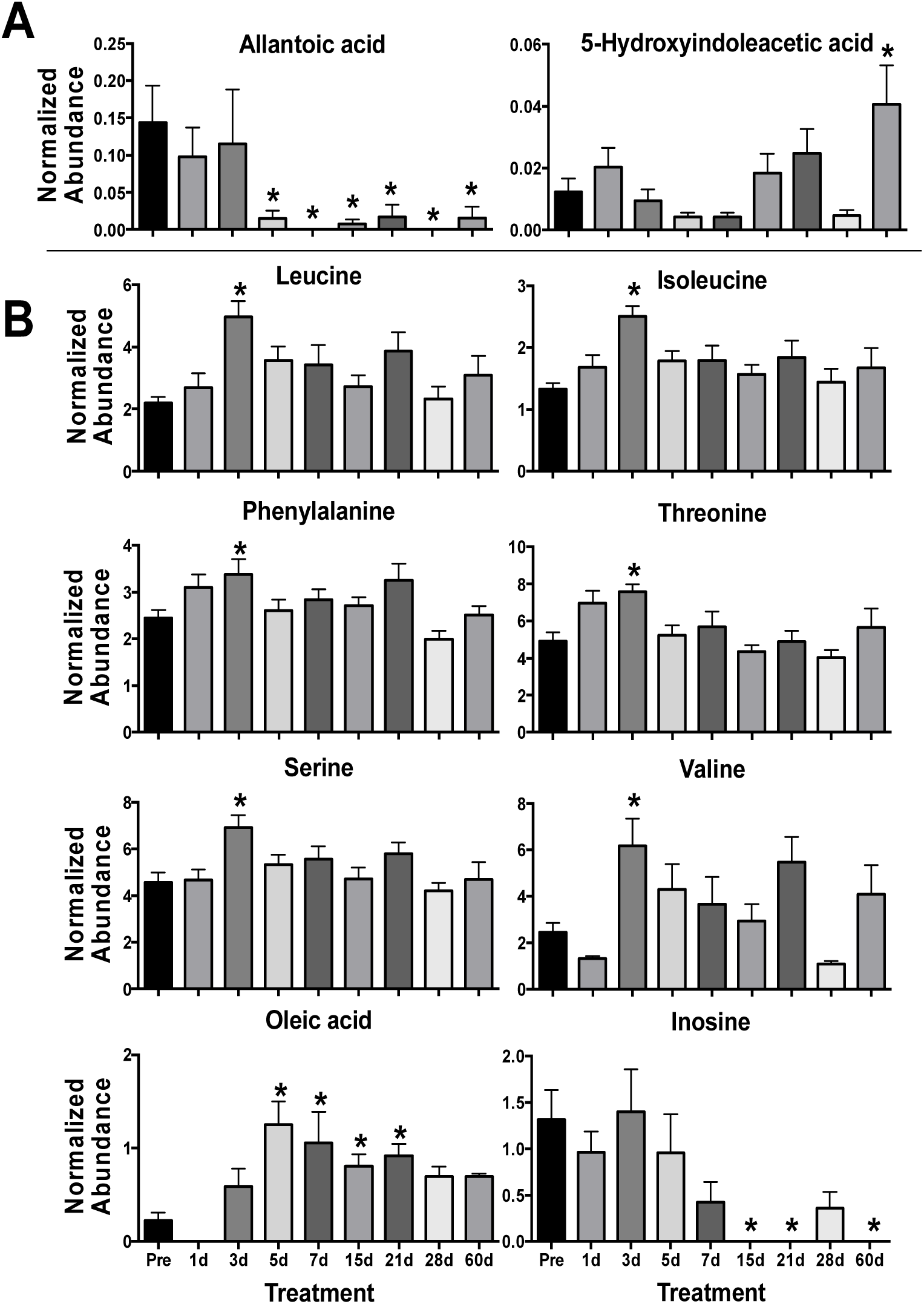
Urine and serum metabolites significantly perturbed after 4 γ Gy TBI in NHPs from 1 – 60 d. (* P < 0.05 determined by a Kruskal-Wallis test and post hoc Dunn’s multiple comparison test, mean ± SEM, pre-exposure samples [-8 and -3 d] were averaged for the control group).

In serum, perturbed levels of several amino acids were evident at 3 d with significantly higher levels of leucine (P = 0.002), isoleucine (P = 0.011), phenylalanine (P = 0.004), threonine (P = 0.003), serine (P = 0.029), and valine (P = 0.003) in the irradiated group (Figure 2B). Amino acids, such as taurine and citrulline, show consistent perturbation due to radiation exposure and are conserved across species [7,40,41]. Our previous work has shown a general trend in decreased NHP serum amino acid levels (primarily in non-essential amino acids) at 7 d at several doses (2, 4, 6, 7, and 10 Gy) that would represent differing ARS severity [20,26,42]. However, fluctuating amino acids levels in the blood over time post-exposure [19,43] clearly portray a much more dynamic pathophysiological response. As a majority of the amino acids identified in this study (except serine) are essential amino acids, diet and nutrient absorption may play a role in observed serum amino acid concentration. Also, sex differences can contribute to basal amino acid levels, such as citrulline [44], and have differential responses after radiation exposure, such as phenylalanine [21]. Here, females showed increased perturbation for phenylalanine (♀ P = 0.027, ♂ P = 0.232), threonine (♀ P < 0.001, ♂ P = 0.251), serine (♀ P = 0.014, ♂ P = 0.123), and valine (♀ P = 0.004, ♂ P = 0.105) at 3 d (Figure 3). As sex differences have been reported in multiple animal models, including NHPs [25,42] and humans [21,45], it must be continually addressed further in biomarker response [46].

**Figure 3.**
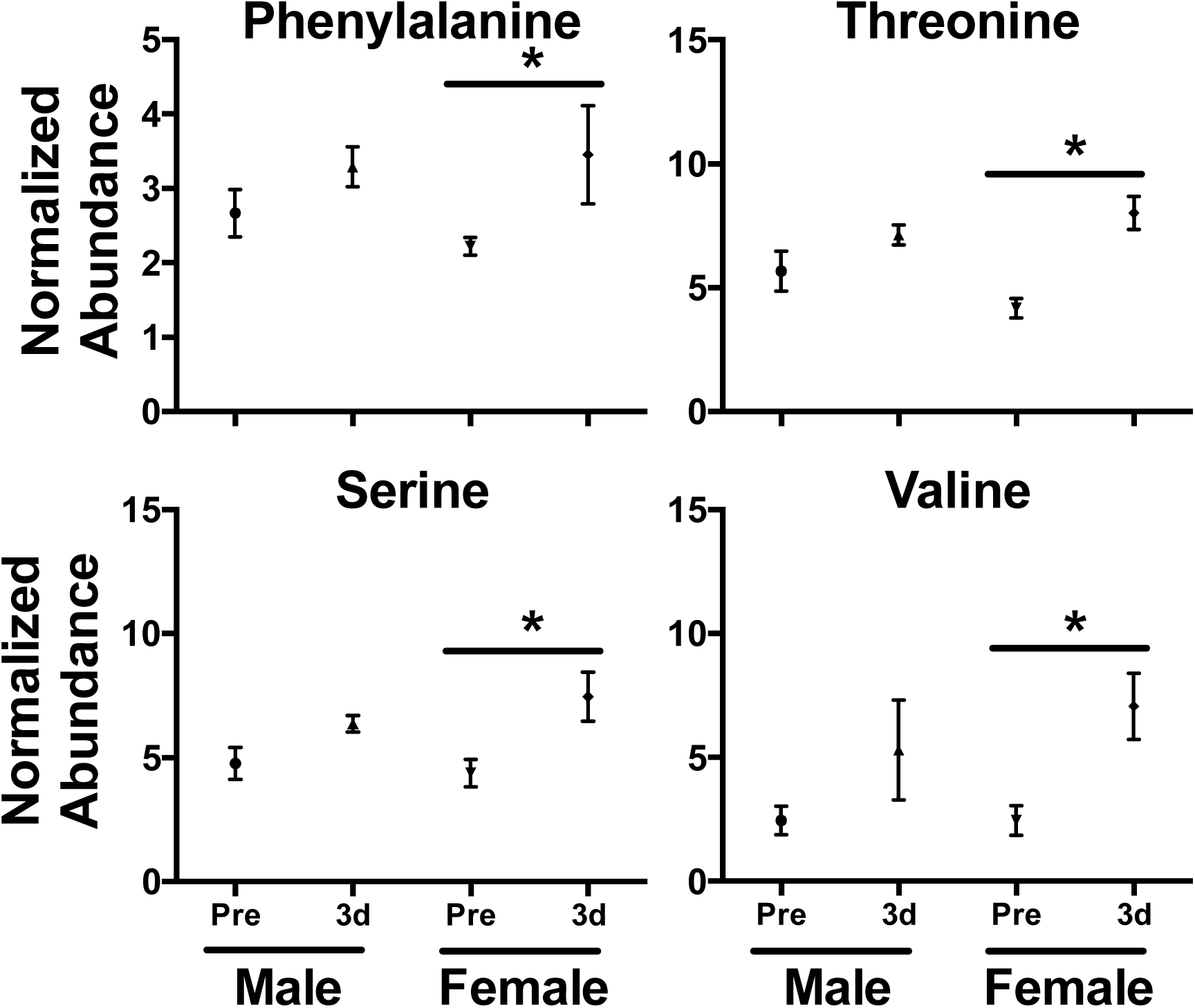
Sex differences in serum amino acid levels 3 d after a 4 Gy γ-ray TBI in NHPs. While levels increased for both males and females after exposure, higher fold changes were observed in phenylalanine, threonine, serine, and valine in females. (* P < 0.05 determined by a t-test, mean ± SEM., pre-exposure samples [-8 and -3 d] were averaged for the control group)

Levels of inosine (P = 0.002) decreased from 15 – 60 d and oleic acid (P < 0.001) increased from 5 – 12 d (Table S1, Figure 2B). Inosine consists of ribose with a C1 linkage to hypoxanthine that can be cleaved by nucleoside phosphorylases. It is an intermediate in purine metabolism and cleavage to hypoxanthine can further lead to downstream metabolites including xanthine and uric acid (again common metabolites detected in radiation metabolomics) and may feed into amino acid metabolism [47,48]. Perturbed blood inosine levels have also been observed in disease such as uremia from kidney dysfunction [49] and sepsis [50], both of which are concerns after potential radiation injury. Oleic acid is a ubiquitous monounsaturated free fatty acid (FFA 18:1) and is less readily oxidized from free radical formation after radiation exposure than other polyunsaturated fatty acid (e.g., linoleic acid or α-linolenic acid) [51]. While sex differences in fatty acids were not observed in this study they have been documented in TBI human patients, where females had slightly higher fold changes in oleic acid and α-linolenic acid compared to males [21]. While not a focus in this study’s approach, the serum lipidome has been the focus of many previous radiation exposure studies and due to the number and complexity of lipid compounds, their importance in cell signaling, and susceptibility to peroxidation make for attractive targets when assessing the indirect cellular effects of IR-induced reactive oxygen species [26,52–55].

In our previous work, we showed significant decreases in urinary TCA metabolites at 7 d after multiple doses of IR in a separate independent cohort [20]. Citric and isocitric acid were significantly lower at all tested doses (2, 4, 6, 7, and 10 Gy) while succinic, *cis*-aconitic, fumaric, and malic acid remained closer to basal levels at 2 and 4 Gy but decreased at ≥ 6 Gy doses. Although some TCA intermediate levels were not significantly decreased at 7 d after 4 Gy, given the importance of the TCA cycle in energy metabolism [56] we used a GC-TQ-MS platform to further explore fold changes in the first week (1, 3, 5, and 7 d) after IR exposure in this separate cohort (irradiated at the same facility > 3 years apart). With the exception of isocitric acid at 1 d, all TCA metabolites showed decreased levels compared to controls (Table 3). Comparing samples at 7 d to the previous cohort, fold changes were similar with decreases in citric and isocitric acid and negligible changes in fumaric and *cis*-aconitic acid, but decreases in malic and succinic acid were slightly greater than previously reported (Table S2). However, when comparing the temporal effects of TBI on urinary TCA cycle intermediates during the first week, the greatest fold changes were seen at 3 d with less pronounced decreases at 1 d (except isocitric acid) and returning closer to baseline levels at 5 – 7 d (Table 3). Urinary citric acid levels have shown a similar trend in mice after an 8 Gy exposure (LD_50/30_ for male C57Bl/6) [57]. Pyruvic acid remained consistently lower from 1 – 7 d. As mitochondria are known sensitive targets of IR induced damage, in part to their overall mass of total cell volume and high reactive oxygen species levels, perturbations to these TCA intermediates may be a direct indication of mitochondrial dysfunction [58].

**Table 3.** Log fold changes (logFC) of urinary pyruvic acid and TCA cycle intermediates at 1, 3, 5, and 7 d after a 4 Gy γ radiation exposure in NHPs.

| Biofluid | Metabolite | HMDB<br>ID | 1 d<br>logFC | 3 d<br>logFC | 5 d<br>logFC | 7 d<br>logFC |
| --- | --- | --- | --- | --- | --- | --- |
| Urine | Pyruvic acid | HMDB 00243 | -0.55 | -0.39 | -0.48 | -0.35 |
|  | Citric acid | HMDB 00094 | -0.02 | -0.58 | -0.24 | -0.27 |
|  | Isocitric acid | HMDB 00193 | 0.12 | -0.49 | -0.25 | -0.20 |
|  | *cis*-Aconitic acid | HMDB 00072 | -0.30 | -0.90 | -0.66 | -0.08 |
| | α-Ketoglutaric acid | HMDB 00208 | -0.17 | -1.02 | -0.65 | -0.10 |
|  | Malic acid | HMDB 00744 | -0.28 | -0.60 | -0.43 | -0.28 |
|  | Succinic acid | HMDB 00254 | -0.24 | -0.79 | -0.38 | -0.21 |
|  | Fumaric acid | HMDB 00134 | -0.27 | -0.70 | -0.57 | -0.11 |

Radiation biodosimetry requires determining biomarkers to serve as a proxy for ARS severity and assignment of appropriate medical treatments. Here, a 4 Gy γ-ray TBI exposure was chosen to limit mortality and allow animal assessment over a 1 – 60 d time course. The LD_50/60_ in this NHPs model is ∼6.6 Gy, which is consistent with other studies conducted without blood transfusion and antibiotics as supportive care [59] and with advanced supportive care (i.e., adding blood transfusion and antibiotics) the LD_50/60_ increase to ∼7.5 Gy [60]. A 4 Gy γ-ray TBI exposure in NHPs is comparable to a ∼2.2 Gy dose in humans where the LD_50/60_ may range from ∼2.5 – 6.0 Gy depending on multiple factors including combined injury, pre-existing conditions, radiosensitivity, and time to supportive care [61,62]. Generally, doses > 2 Gy in humans will cause ARS associated hematopoietic and require medical care; conversely, little to any survival is expected at doses > 10 Gy [61]. In the event of an accidental or intentional radiation exposure, it is unlikely individuals will reach emergency response stations within 24 h, which necessitates relevant markers in the 1 – 7 d range allowing realistic time for triage. At 15 – 60 d symptoms of hematopoietic syndrome (e.g., thrombocytopenia and neutropenia) will progress to more advanced stages but may still require medical intervention, such as hematopoietic cell transplantation in severe cases, and observation for longer-term effects. These results show that metabolomic signatures can elucidate phenotypic changes, such as mitochondrial dysfunction and amino acid perturbation, allowing differentiation among pre-exposure and post-exposure groups and possible longer-term effects from radiation exposure.

## 4. Conclusions

The majority of studies defining biophysiological responses to radiation exposures have focused on LC-MS platforms [7], however, these methods lack in chromatographic separation of small volatile compounds or isomers (e.g., leucine vs. isoleucine) and availability of well-established RI and mass spectral libraries available in the realm of GC-MS analyses. While longer run times (∼30 min vs. ∼5-10 min) and sample preparation for required derivatization (∼60 min vs. ∼5 min) limits its direct use in biodosimetry for time-sensitive emergency situations, GC-MS serves as a complementary platform to LC-MS providing more depth into metabolite coverage, such as the TCA cycle intermediates described in this study. Subsequent compound analyses can be further refined for high-throughput targeted platforms more suited to biodosimetry, such as differential mobility spectrometry (DMS) MS [63,64] or LC-MS [24]. In this study, we examined global small molecule signatures in NHP biofluids (urine and serum) from 1 – 60 d after a 4 Gy γ-ray TBI exposure and determined fold changes in urinary TCA cycle intermediates from 1 – 7 d using GC-MS platforms. We found perturbed metabolites in NHP biofluids involved in amino acid, lipid, serotonin, and purine metabolism, some of which may reflect dietary and host microbiome changes. Primarily essential amino acids were identified in this study and females exhibited higher fold changes (including non-essential serine), but influences of diet and nutrient absorption on essential amino acid levels should be examined if used as a proxy for radiation injury without additional multivariate analyses [43,65]. Finally, we found the greatest perturbation of TCA intermediates in the first week at ∼3 d and this trend may be consistent across species [57]. These results highlight the importance of using multiple analytical platforms when examining biophysiological responses to radiation injury and future work may integrate these results into a single platform for rapid biodosimetry.

## Author Contributions

conceptualization, E.L.P., E.C.L., and A.J.F.; methodology (mass spectrometry), E.L.P., M.G., S.E.D., and S.D.; formal analysis, E.L.P., E.C.L., M.G., and S.E.D.; investigation, E.L.P., E.C.L., M.G., S.E.D., and A.J.F.; resources, S.D., D.N., K.B., J.B., J.G., S.A., P.C., and A.J.F.; data curation, E.L.P., E.C.L., M.G., S.E.D., and S.D.; writing—original draft preparation, E.L.P., E.C.L., and A.J.F.; writing—review and editing, E.L.P., E.C.L., M.G., S.E.D., S.D., D.N., K.B., J.B., J.G., S.A., P.C., and A.J.F.; visualization, E.L.P., E.C.L., and A.J.F.; supervision, E.L.P., E.C.L., and A.J.F.; project administration, E.C.L., S.D., D.N., K.B., J.B., J.G., S.A., P.C., and A.J.F.; funding acquisition, A.J.F.

## Funding and Acknowledgments

This work was funded by a pilot grant (P.I. ELP) from the Opportunity Funds Management Core of the Centers for Medical Countermeasures against Radiation, National Institute of Allergy and Infectious Diseases (NIAID) (grant # U19AI067773; P.I. David Brenner), under HHS Contract (HHSN272201500013I) awarded to SRI International and NIAID grant # 1RO1AI101798 (P.I. AJF). The authors acknowledge Lombardi Comprehensive Cancer Metabolomics Shared Resource (MSR) for help with data acquisition, which has partial support from National Cancer Institute grant # P30CA051008 (P.I. Louis Weiner). Content is the responsibility of authors and does not necessarily represent official views of NIH.

---

**Table S1.** Normalized abundance of compounds detected by GC-TOF-MS global profiling of NHP biofluids after 4 Gy γ radiation exposure.

|  | **Metabolite name** | **Pre** | **1 d** | **3 d** | **5 d** | **7 d** | **15 d** | **21 d** | **28 d** | **60 d** |
| --- | --- | --- | --- | --- | --- | --- | --- | --- | --- | --- |
| Urine | Allantoic acid | 0.14±0.05 | 0.10±0.04 | 0.12±0.07 | 0.02±0.01 | 0.00±0.00 | 0.01±0.01 | 0.02±0.02 | 0.00±0.00 | 0.02±0.02 |
|  | 5-Hydroxyindoleacetic acid | 0.01±0.00 | 0.02±0.01 | 0.01±0.00 | 0.00±0.00 | 0.00±0.00 | 0.02±0.01 | 0.02±0.01 | 0.00±0.00 | 0.04±0.01 |
| Serum | Oleic acid | 0.23±0.09 | 0.00±0.00 | 0.59±0.19 | 1.25±0.25 | 1.06±0.33 | 0.81±0.13 | 0.92±0.13 | 0.69±0.11 | 0.69±0.03 |
|  | Inosine | 1.31±0.32 | 0.96±0.22 | 1.40±0.46 | 0.96±0.41 | 0.42±0.22 | 0.00±0.00 | 0.00±0.00 | 0.36±0.18 | 0.00±0.00 |
|  | Leucine | 2.20±0.20 | 2.70±0.46 | 4.97±0.51 | 3.57±0.44 | 3.43±0.63 | 2.73±0.36 | 3.86±0.61 | 2.32±0.40 | 3.09±0.64 |
|  | Isoleucine | 1.33±0.09 | 1.68±0.20 | 2.51±0.17 | 1.78±0.16 | 1.80±0.24 | 1.57±0.15 | 1.85±0.27 | 1.44±0.21 | 1.67±0.32 |
|  | Valine | 0.23±0.09 | 0.00±0.00 | 0.59±0.19 | 1.25±0.25 | 1.06±0.33 | 0.81±0.13 | 0.92±0.13 | 0.69±0.11 | 0.69±0.03 |
|  | Serine | 4.57±0.41 | 4.69±0.43 | 6.92±0.53 | 5.32±0.44 | 5.57±0.55 | 4.71±0.49 | 5.81±0.46 | 4.20±0.36 | 4.71±0.73 |
|  | Threonine | 4.92±0.48 | 6.98±0.67 | 7.58±0.40 | 5.23±0.54 | 5.71±0.83 | 4.36±0.35 | 4.90±0.57 | 4.03±0.41 | 5.66±1.02 |
|  | Phenylalanine | 2.45±0.17 | 3.10±0.27 | 3.38±0.33 | 2.60±0.24 | 2.83±0.23 | 2.72±0.17 | 3.26±0.36 | 1.99±0.18 | 2.51±0.19 |

(Mean ± SEM)

**Table S2.** Comparisons of log fold changes (logFC) of urinary TCA cycle intermediates at 7 d after a 4 Gy γ radiation exposure in a previous NHP cohort [25,26] and the current cohort.

| **Biofluid** | **Metabolite** | **logFC Control vs. 7 d**<br>**Previous Cohort** | **logFC Control vs. 7 d**<br>**Current Cohort** |
| --- | --- | --- | --- |
| Urine | Citric acid | -0.35 | -0.27 |
|  | Isocitric acid | -0.26 | -0.20 |
|  | *cis*-Aconitic acid | -0.18 | -0.08 |
|  | Malic acid | -0.11 | -0.26 |
|  | Succinic acid | -0.13 | -0.21 |
|  | Fumaric acid | 0.05 | -0.11 |

## References

1. Czarwinski, R.; Crick, M.J. Occupational exposures worldwide and revision of international standards for protection. Radiat Prot Dosimetry 2011, 144, 2–11.

2. Izrael, Y.A. Chernobyl radionuclide distribution and migration. Health Phys 2007, 93, 410–417.

3. McLaughlin, J.P. Some characteristics and effects of natural radiation. Radiat Prot Dosimetry 2015, 167, 2–7.

4. Ozasa, K.; Grant, E.J.; Kodama, K. Japanese legacy cohorts: The life span study atomic bomb survivor cohort and survivors’ offspring. J Epidemiol 2018, 28, 162–169.

5. Ozasa, K.; Cullings, H.M.; Ohishi, W.; Hida, A.; Grant, E.J. Epidemiological studies of atomic bomb radiation at the Radiation Effects Research Foundation. Int J Radiat Biol 2019, 1-13.

6. DiCarlo, A.L.; Ramakrishnan, N.; Hatchett, R.J. Radiation combined injury: Overview of NIAID research. Health Phys 2010, 98, 863–867.

7. Pannkuk, E.L.; Fornace Jr, A.J.; Laiakis, E.C. Metabolomic applications in radiation biodosimetry: Exploring radiation effects through small molecules. Int J Radiat Biol 2017, 93, 1151–1176.

8. Johnson, C.H.; Ivanisevic, J.; Siuzdak, G. Metabolomics: Beyond biomarkers and towards mechanisms. Nat Rev Mol Cell Biol 2016, 17, 451–459.

9. Gowda, G.A.; Djukovic, D. Overview of mass spectrometry-based metabolomics: opportunities and challenges. Methods Mol Biol 2014, 1198, 3–12.

10. Markley, J.L.; Brüschweiler, R.; Edison, A.S.; Eghbalnia, H.R.; Powers, R.; Raftery, D.; Wishart, D.S. The future of NMR-based metabolomics. Curr Opin Biotechnol 2017, 43, 34–40.

11. Fuhrer, T.; Zamboni, N. High-throughput discovery metabolomics. Curr Opin Biotechnol 2015, 31, 73–78.

12. Horning, E.C.; Horning, M.G. Metabolic profiles: Chromatographic methods for isolation and characterization of a variety of metabolites in man. Methods Med Res 1970, 12, 369–371.

13. Horning, E.C.; Horning, M.G. Metabolic profiles: Gas-phase methods for analysis of metabolites. Clin Chem 1971, 17, 802–809.

14. Fiehn, O. Metabolomics by gas chromatography-mass spectrometry: Combined targeted and untargeted profiling. Curr Protoc Mol Biol 2016, 114, 30.4.1-30.4.32.

15. Beale, D.J.; Pinu, F.R.; Kouremenos, K.A.; Poojary, M.M.; Narayana, V.K.; Boughton, B.A.; Kanojia, K.; Dayalan, S.; Jones, O.A.H.; Dias, D.A. Review of recent developments in GC-MS approaches to metabolomics-based research. Metabolomics 2018, 14, 152–152.

16. Lee, D.Y.; Bowen, B.P.; Nguyen, D.H.; Parsa, S.; Huang, Y.; Mao, J.H.; Northen, T.R. Low-dose ionizing radiation-induced blood plasma metabolic response in a diverse genetic mouse population. Radiat Res 2012, 178, 551–555.

17. Lanz, C.; Patterson, A.D.; Slavik, J.; Krausz, K.W.; Ledermann, M.; Gonzalez, F.J.; Idle, J.R. Radiation metabolomics. 3. Biomarker discovery in the urine of gamma-irradiated rats using a simplified metabolomics protocol of gas chromatography-mass spectrometry combined with random forests machine learning algorithm. Radiat Res 2009, 172, 198–212.

18. Zhao, M.; Lau, K.K.; Zhou, X.; Wu, J.; Yang, J.; Wang, C. Urinary metabolic signatures and early triage of acute radiation exposure in rat model. Mol Biosyst 2017, 13, 756–766.

19. Liu, H.; Wang, Z.; Zhang, X.; Qiao, Y.; Wu, S.; Dong, F.; Chen, Y. Selection of candidate radiation biomarkers in the serum of rats exposed to gamma-rays by GC/TOFMS-based metabolomics. Radiat Prot Dosimetry 2013, 154, 9–17.

20. Pannkuk, E.L.; Laiakis, E.C.; Authier, S.; Wong, K.; Fornace Jr, A.J. Gas chromatography/mass spectrometry metabolomics of urine and serum from nonhuman primates exposed to ionizing radiation: Impacts on the tricarboxylic acid cycle and protein metabolism. J Proteome Res 2017, 16, 2091–2100.

21. Laiakis, E.C.; Pannkuk, E.L.; Chauthe, S.K.; Wang, Y.W.; Lian, M.; Mak, T.D.; Barker, C.A.; Astarita, G.; Fornace Jr, A.J. A serum small molecule biosignature of radiation exposure from total body irradiated patients. J Proteome Res 2017, 16, 3805–3815.

22. Mörén, L.; Wibom, C.; Bergström, P.; Johansson, M.; Antti, H.; Bergenheim, A.T. Characterization of the serum metabolome following radiation treatment in patients with high-grade gliomas. Radiat Oncol 2016, 11, 51.

23. Singh, V.K.; Newman, V.L.; Berg, A.N.; MacVittie, T.J. Animal models for acute radiation syndrome drug discovery. Expert Opin Drug Discov 2015, 10, 497–517.

24. Pannkuk, E.L.; Laiakis, E.C.; Gill, K.; Jain, S.; Mehta, K.; Nishita, D.; Bujold, K.; Bakke, J.; Gahagen, J.; Authier, S.; Chang, P.; Fornace Jr, A.J. Liquid chromatography - mass spectrometry based metabolomics of nonhuman primates after 4 Gy total body radiation exposure: Global effects and targeted panels. Journal of Proteome Research 2019,

25. Pannkuk, E.L.; Laiakis, E.C.; Authier, S.; Wong, K.; Fornace Jr, A.J. Global metabolomic identification of longer-term dose dependent urinary biomarkers in nonhuman primates exposed to ionizing radiation Radiat Res 2015, 184, 121–131.

26. Pannkuk, E.L.; Laiakis, E.C.; Mak, T.D.; Astarita, G.; Authier, S.; Wong, K.; Fornace Jr, A.J. A lipidomic and metabolomic serum signature from nonhuman primates exposed to ionizing radiation Metabolomics 2016, 12, 1–11.

27. Chong, J.; Soufan, O.; Li, C.; Caraus, I.; Li, S.; Bourque, G.; Wishart, D.S.; Xia, J. MetaboAnalyst 4.0: Towards more transparent and integrative metabolomics analysis. Nucleic Acids Res 2018, 46, W486–W494.

28. Bijlsma, S.; Bobeldijk, I.; Verheij, E.R.; Ramaker, R.; Kochhar, S.; Macdonald, I.A.; van Ommen, B.; Smilde, A.K. Large-scale human metabolomics studies: A strategy for data (pre-) processing and validation. Anal Chem 2006, 78, 567–574.

29. Ramazzina, I.; Folli, C.; Secchi, A.; Berni, R.; Percudani, R. Completing the uric acid degradation pathway through phylogenetic comparison of whole genomes. Nat Chem Biol 2006, 2, 144–148.

30. Usuda, N.; Hayashi, S.; Fujiwara, S.; Noguchi, T.; Nagata, T.; Rao, M.S.; Alvares, K.; Reddy, J.K.; Yeldandi, A.V. Uric acid degrading enzymes, urate oxidase and allantoinase, are associated with different subcellular organelles in frog liver and kidney. J Cell Sci 1994, 107, 1073–1081.

31. Kratzer, J.T.; Lanaspa, M.A.; Murphy, M.N.; Cicerchi, C.; Graves, C.L.; Tipton, P.A.; Ortlund, E.A.; Johnson, R.J.; Gaucher, E.A. Evolutionary history and metabolic insights of ancient mammalian uricases. Proc Natl Acad Sci U S A 2014, 111, 3763–3768.

32. Yano, J.M.; Yu, K.; Donaldson, G.P.; Shastri, G.G.; Ann, P.; Ma, L.; Nagler, C.R.; Ismagilov, R.F.; Mazmanian, S.K.; Hsiao, E.Y. Indigenous bacteria from the gut microbiota regulate host serotonin biosynthesis. Cell 2015, 161, 264–276.

33. Christenson, J.G.; Dairman, W.; Udenfriend, S. On the identity of DOPA decarboxylase and 5-hydroxytryptophan decarboxylase (immunological titration-aromatic L-amino acid decarboxylase-serotonin-dopamine-norepinephrine). Proc Natl Acad Sci U S A 1972, 69, 343–347.

34. Kobayashi, S.; Nakamura, W.; Eto, H. Protective effect of 5-hydroxytryptophan against lethal doses of x-radiation in mice. Int J Radiat Biol Relat Stud Phys Chem Med 1966, 11, 505–508.

35. Barnes, J.H.; Lowman, D.M. Relative radioprotective abilities of 5-hydroxytryptophan and 5-hydroxytryptamine. Int J Radiat Biol Relat Stud Phys Chem Med 1968, 14, 87–88.

36. Randic, M.; Supek, Z. Urinary excretion of 5-hydroxyindolacetic acid after a single whole-body x-irradiation in normal and adrenalectomized rats. Int J Radiat Biol Relat Stud Phys Chem Med 1961, 4, 151–153.

37. Grison, S.; Favé, G.; Maillot, M.; Manens, L.; Delissen, O.; Blanchardon, E.; Banzet, N.; Defoort, C.; Bott, R.; Dublineau, I.; Aigueperse, J.; Gourmelon, P.; Martin, J.C.; Souidi, M. Metabolomics identifies a biological response to chronic low-dose natural uranium contamination in urine samples. Metabolomics 2013, 9, 1168–1180.

38. Deanovic, Z.; Supek, Z.; Randic, M. Relationship between the dose of whole-body x-irradiation and the urinary excretion of 5-hydroxyindoleacetic acid in rats. Int J Radiat Biol Relat Stud Phys Chem Med 1963, 7, 569–574.

39. ÓBroin, P.; Vaitheesvaran, B.; Saha, S.; Hartil, K.; Chen, E.I.; Goldman, D.; Fleming, W.H.; Kurland, I.J.; Guha, C.; Golden, A. Intestinal microbiota-derived metabolomic blood plasma markers for prior radiation injury. Int J Radiat Oncol Biol Phys 2015, 91, 360–367.

40. Johnson, C.H.; Patterson, A.D.; Krausz, K.W.; Kalinich, J.F.; Tyburski, J.B.; Kang, D.W.; Luecke, H.; Gonzalez, F.J.; Blakely, W.F.; Idle, J.R. Radiation metabolomics. 5. Identification of urinary biomarkers of ionizing radiation exposure in nonhuman primates by mass spectrometry-based metabolomics. Radiat Res 2012, 178, 328–340.

41. Jones, J.W.; Tudor, G.; Bennett, A.; Farese, A.M.; Moroni, M.; Booth, C.; MacVittie, T.J.; Kane, M.A. Development and validation of a LC-MS/MS assay for quantitation of plasma citrulline for application to animal models of the acute radiation syndrome across multiple species. Anal Bioanal Chem 2014, 406, 4663–4675.

42. Pannkuk, E.L.; Laiakis, E.C.; Authier, S.; Wong, K.; Fornace Jr, A.J. Targeted metabolomics of nonhuman primate serum after exposure to ionizing radiation: Potential tools for highthroughput biodosimetry. RSC Advances 2016, 6, 51192–51202.

43. Tang, X.; Zheng, M.; Zhang, Y.; Fan, S.; Wang, C. Estimation value of plasma amino acid target analysis to the acute radiation injury early triage in the rat model. Metabolomics 2013, 9, 853–863.

44. Bujold, K.; Hauer-Jensen, M.; Donini, O.; Rumage, A.; Hartman, D.; Hendrickson, H.P.; Stamatopoulos, J.; Naraghi, H.; Pouliot, M.; Ascah, A.; Sebastian, M.; Pugsley, M.K.; Wong, K.; Authier, S. Citrulline as a biomarker for gastrointestinal-acute radiation syndrome: Species differences and experimental condition effects. Radiat Res 2016, 186, 71–78.

45. Laiakis, E.C.; Mak, T.D.; Anizan, S.; Amundson, S.A.; Barker, C.A.; Wolden, S.L.; Brenner, D.J.; Fornace Jr, A.J. Development of a metabolomic radiation signature in urine from patients undergoing total body irradiation. Radiat Res 2014, 181, 350–361.

46. Jones, J.W.; Alloush, J.; Sellamuthu, R.; Chua, H.L.; MacVittie, T.J.; Orschell, C.M.; Kane, M.A. Effect of sex on biomarker response in a mouse model of the hematopoietic acute radiation syndrome. Health Phys 2019, 116, 484–502.

47. Simoni, R.E.; Gomes, L.N.; Scalco, F.B.; Oliveira, C.P.; Aquino Neto, F.R.; de Oliveira, M.L. Uric acid changes in urine and plasma: An effective tool in screening for purine inborn errors of metabolism and other pathological conditions. J Inherit Metab Dis 2007, 30, 295–309.

48. Tyburski, J.B.; Patterson, A.D.; Krausz, K.W.; Slavik, J.; Fornace Jr, A.J.; Gonzalez, F.J.; Idle, J.R. Radiation metabolomics. 2. Dose- and time-dependent urinary excretion of deaminated purines and pyrimidines after sublethal gamma-radiation exposure in mice. Radiat Res 2009, 172, 42–57.

49. Niwa, T.; Takeda, N.; Yoshizumi, H. RNA metabolism in uremic patients: Accumulation of modified ribonucleosides in uremic serum. Technical note. Kidney Int 1998, 53, 1801–1806.

50. Jabs, C.M.; Sigurdsson, G.H.; Neglen, P. Plasma levels of high-energy compounds compared with severity of illness in critically ill patients in the intensive care unit. Surgery 1998, 124, 65–72.

51. Yin, H.; Xu, L.; Porter, N.A. Free radical lipid peroxidation: Mechanisms and analysis. Chem Rev 2011, 111, 5944–5972.

52. Tyurina, Y.Y.; Tyurin, V.A.; Kapralova, V.I.; Wasserloos, K.; Mosher, M.; Epperly, M.W.; Greenberger, J.S.; Pitt, B.R.; Kagan, V.E. Oxidative lipidomics of γ-radiation-induced lung injury: Mass spectrometric characterization of cardiolipin and phosphatidylserine peroxidation. Radiat Res 2011, 175, 610–621.

53. Carter, C.L.; Jones, J.W.; Barrow, K.; Kieta, K.; Taylor-Howell, C.; Kearney, S.; Smith, C.P.; Gibbs, A.; Farese, A.M.; MacVittie, T.J.; Kane, M.A. A MALDI-MSI approach to the characterization of radiation-induced lung injury and medical countermeasure development. Health Phys 2015, 109, 466–478.

54. Goudarzi, M.; Weber, W.M.; Chung, J.; Doyle-Eisele, M.; Melo, D.R.; Mak, T.D.; Strawn, S.J.; Brenner, D.J.; Guilmette, R.; Fornace Jr, A.J. Serum dyslipidemia is induced by internal exposure to strontium-90 in mice, lipidomic profiling using a data-independent liquid chromatography-mass spectrometry approach. J Proteome Res 2015, 14, 4039–4049.

55. Laiakis, E.C.; Strassburg, K.; Bogumil, R.; Lai, S.; Vreeken, R.J.; Hankemeier, T.; Langridge, J.; Plumb, R.S.; Fornace Jr, A.J.; Astarita, G. Metabolic phenotyping reveals a lipid mediator response to ionizing radiation. J Proteome Res 2014, 13, 4143–4154.

56. Williamson, J.R.; Cooper, R.H. Regulation of the citric acid cycle in mammalian systems. FEBS Lett 1980, 117 Suppl, K73-85.

57. Chen, C.; Brenner, D.J.; Brown, T.R. Identification of urinary biomarkers from x-irradiated mice using NMR spectroscopy. Radiat Res 2011, 175, 622–630.

58. Azzam, E.I.; Jay-Gerin, J.P.; Pain, D. Ionizing radiation-induced metabolic oxidative stress and prolonged cell injury. Cancer Lett 2012, 327, 48–60.

59. Singh, V.K.; Kulkarni, S.; Fatanmi, O.O.; Wise, S.Y.; Newman, V.L.; Romaine, P.L.; Hendrickson, H.; Gulani, J.; Ghosh, S.P.; Kumar, K.S.; Hauer-Jensen, M. Radioprotective efficacy of gamma-tocotrienol in nonhuman primates. Radiat Res 2016, 185, 285–298.

60. Farese, A.M.; Cohen, M.V.; Katz, B.P.; Smith, C.P.; Jackson, W.; Cohen, D.M.; MacVittie, T.J. A nonhuman primate model of the hematopoietic acute radiation syndrome plus medical management. Health Phys 2012, 103, 367–382.

61. López, M.; Martín, M. Medical management of the acute radiation syndrome. Rep Pract Oncol Radiother 2011, 16, 138–146.

62. Macià I Garau, M.; Lucas Calduch, A.; López, E.C. Radiobiology of the acute radiation syndrome. Rep Pract Oncol Radiother 2011, 16, 123–130.

63. Vera, N.B.; Chen, Z.; Pannkuk, E.L.; Laiakis, E.C.; Fornace Jr, A.J.; Erion, D.M.; Coy, S.L.; Pfefferkorn, J.A.; Vouros, P. Differential mobility spectrometry (DMS) reveals the elevation of urinary acetylcarnitine in non-human primates (NHPs) exposed to radiation. J Mass Spectrom 2018, 53, 548–559.

64. Chen, Z.; Coy, S.L.; Pannkuk, E.L.; Laiakis, E.C.; Fornace Jr, A.J.; Vouros, P. Differential mobility spectrometry-mass spectrometry (DMS-MS) in radiation biodosimetry: Rapid and high-throughput quantitation of multiple radiation biomarkers in nonhuman primate urine. Journal of the American Society for Mass Spectrometry 2018, 29, 1650–1664.

65. Zhang, Y.; Zhou, X.; Li, C.; Wu, J.; Kuo, J.E.; Wang, C. Assessment of early triage for acute radiation injury in rat model based on urinary amino acid target analysis. Mol Biosyst 2014, 10, 1441–1449.

